# Free energy calculations of protein-water complexes with Gromacs

**DOI:** 10.1101/305037

**Authors:** M. A. Shahzad

## Abstract

We used GFP(Green Fluorescent Protein) to understand its basic structure, adding solvent water around the GFP, minimize and equilibrating it using molecular dynamics simulation with Gromacs. Gromacs is an open source software and widely used in molecular dynamics simulation of biological molecules such as proteins, and nucleic acids (DNA AND RNA-molecules). We employ the CHARMM (Chemistry at HARvard Molecular Mechanics) program for the force fields which enable the potential energy of a molecular system to be calculated rapidly. In this particular simple molecular mechanics interaction fields (CHARMM), the force fields consists of stretching energy, bending energy and torsion energy. The non-bond interaction energy is modeled by Lennard-Jones potential. The `pdb2gmx` gromacs command is implemented to obtained the basic coordinate file and topology for the particular system from the GFP PDB file (1gfl.pdb). We enclosed water molecule in a rhombic dodecahedron box having size 0.5*nm*, and protein are embedded in solvent water. The steepest descent method (first-order minimization) are implemented to calculate the local energy minimum with 10^4^-steps. The stability of protein with solvent water molecule are analyzed by measuring the root-mean square displacement (RMSD) of all atoms. It was shown with help of figure that RMSD initially increases rapidly in the first part of simulation, but become stable around 0.14nm, roughly the resolution of the X-ray structure. The difference is partly due to the moving and vibration of atoms around an equilibrium structure. Secondary structure of GFP protein is presented with the help of DSSP-program.

## I. INTRODUCTION

Gromacs is an engine to perform molecular dynamics simulations and energy minimization of biomolecular systems which usually consists of several tens to thousands of amino acid residues. Molecular dynamics simulation presents the time evolution of a molecular systems such as protein, by numerically solving Newton's equation of motion for all atoms in the system. Molecular dynamics (MD) is a powerful tool which can be used to simulate the time evolution of molecular systems with great accuracy. Gromacs is one of application which able to do molecular dynamics simulation based on equation of Newton's law [1–3]. The algorithm used for molecular dynamics assigns positions and velocities to every atom in the simulation system. The forces are computed, which can be used to evolve the positions and velocities through Newtons law, using a given finite time step. The non-bonded interactions between particles model behavior like van der Waals forces, or Coulombs law.

Here we used molecular dynamics simulation on protein with water solvent. We minimize and equlibirate the system utilizing energy minimization technique with Gromacs. The stability of the system is characterized by studying the RMSD and secondary structure of protein.

## II. RESULT

We used Green fluorescent protein (GFP)-water complexes in our simulation. The Green fluorescent protein (GFP) [4–7] is an important proteins currently used as a marker of gene expression and protein localization, and as a bio-sensor [8]. The green fluorescent protein (GFP) is composed of 238 amino acid residues and a molecular weight of 26.9 kdalton that exhibits bright green fluorescence when exposed to light [9].

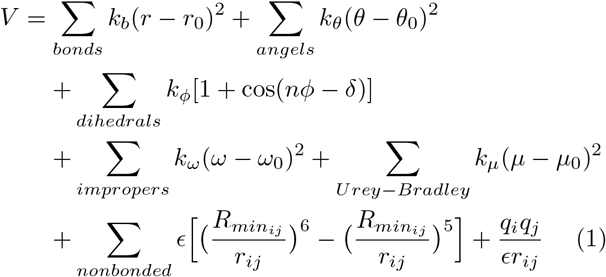

The first term accounts for the bond stretches, with force constant *k_b_* and *r — r_0_* is the distance from equilibrium. The second term in the equation accounts for the bond angles where *k_θ_* is the angle force constant The third term is for the torsion angles where *k_φ_* is the dihedral force constant. The fourth term accounts for the improper, that is out of plane bending, where *k_ω_* is the force constant and *ω — ω*_0_ is the out of plane angle.

The native structure of a GFP is taken from the protein data bank (PDB ID:1gfl) file and the interaction between the amino acids are grouped into native and non-native state. The protein data bank (PDB) format file contains information about atomic coordinates for various types of proteins, small molecules, ions and water. It provides a standard representation for macromolecular structure data which is derived from X-ray diffraction and nuclear magnetic resonance (NMR) studies.

We used pdf2gmx command in Gromacs to convert the pdb (GFP pdb:1gfl.pdb) file format into Gromacs coordinate format, and adds water around GFT. We choose a CHARMM-force fields, given by the following equation [10, 11]:

The Urey-Bradley component (cross-term accounting for angle bending using 1,3 nonbonded interactions) comprises the fifth term, where *k_U_* is the respective force constant and *U* is the distance between the 1,3 atoms in the harmonic potential. Nonbonded interactions between pairs of atoms (*i,j*) are represented by standard 12-6 Lennard-Jones potential and the electrostatic energy with a Coulombic potential. We used periodic boundary conditions for the protein to be simulated to keep track of the motion of all particles We define a rhombic dodecahedron box of size 0.5*nm* To mimic the physiological environment for a protein, we add water molecule around the protein. Fig. (1) illustrate the water molecules around the protein in a box having size 0.5*nm*.

**FIG. 1.**
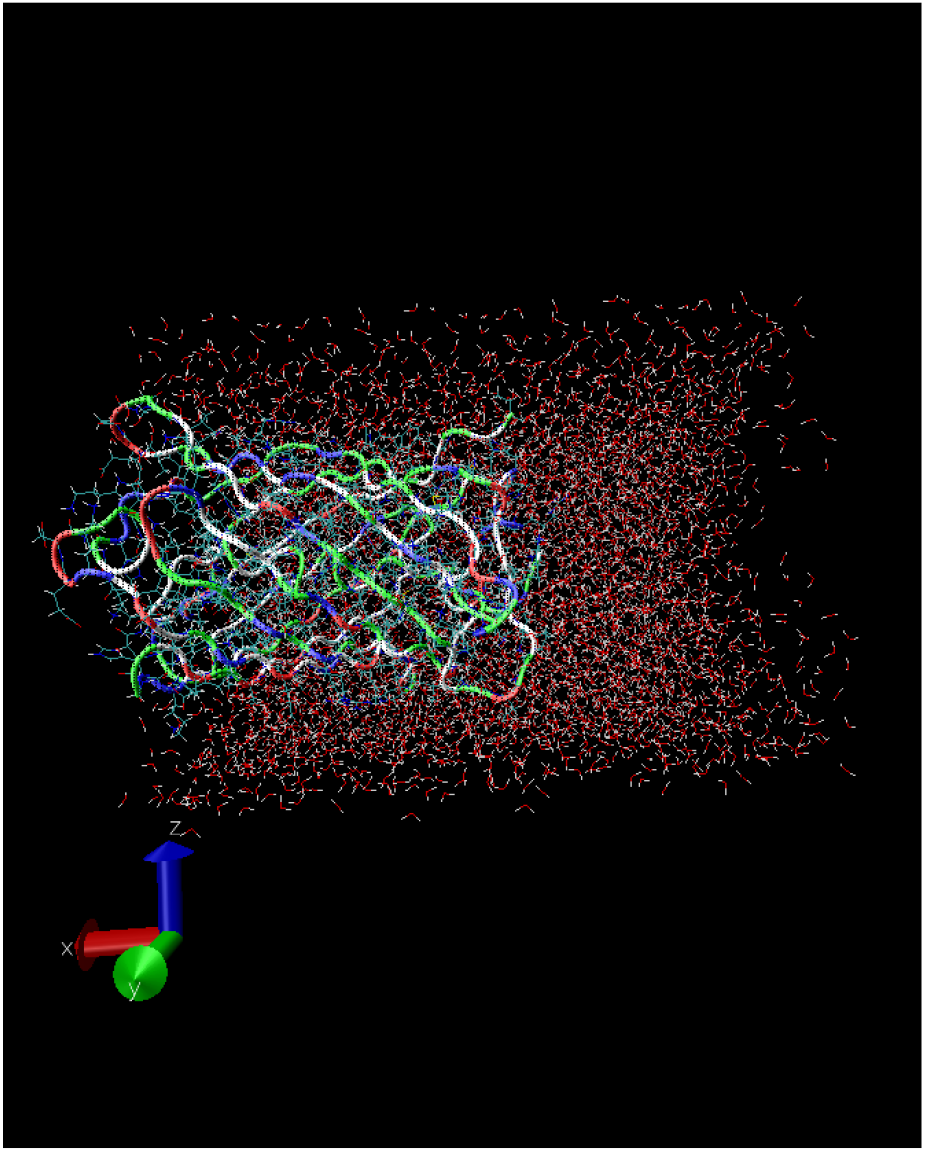
GFP, with colors that vary according to the residues type, embedded in water molecule. Figure produced by VMD-software.

We used md-integrator, a leaf-frog algorithm for integrating Newton's equations of motion. Gromacs also provide a lead-frog stochastic dynamics integrator (sd), and Euler integrator for Langevin dynamics (bd). The leapfrog algorithm implemented in Gromacs uses position **r** and velocity vector **v** at time *t* and *t* — Δ*t*/2, respectively [12–14]. Mathematically:

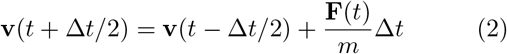

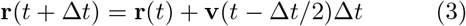

where **F**(*t*) represent the force as a function of time *t*. In our simulation, we used nsteps=10^4^, maximum numbers of steps to integrate. The time steps for integration is dt=0.002. The cut-off distance and neighbor list are rlist=1.0 and nstlist=10, respectively.

To understand the stability of the constructed system used as starting structure for the molecular dynamic simulations, the `Protein-H` root mean square deviation (RMSD) with respect to the `Protein-H` group were calculated. The RMSD in Gromcas can be calculated by using the formula,

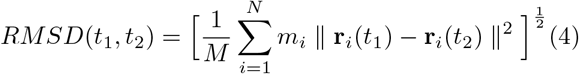

where **r**_*i*_(t) is the position of *i*th atom at time *t*, and *M* = Σ *m_i_* represent the total mass. Fig. (2), represents the root mean square deviation (RMSD) of GFP-water complex over the equilibration time of 200*ps*. Initially the RMSD increase rapidly, but stabilizes around 0.14*nm*. To see that the system (GFP-water) is at an energy minimum, we plot the average potential energy *E_pot_* as a function of energy minimization steps. Fig. (3) illustrate stability and convergence of the potential energy.

**FIG. 2.**
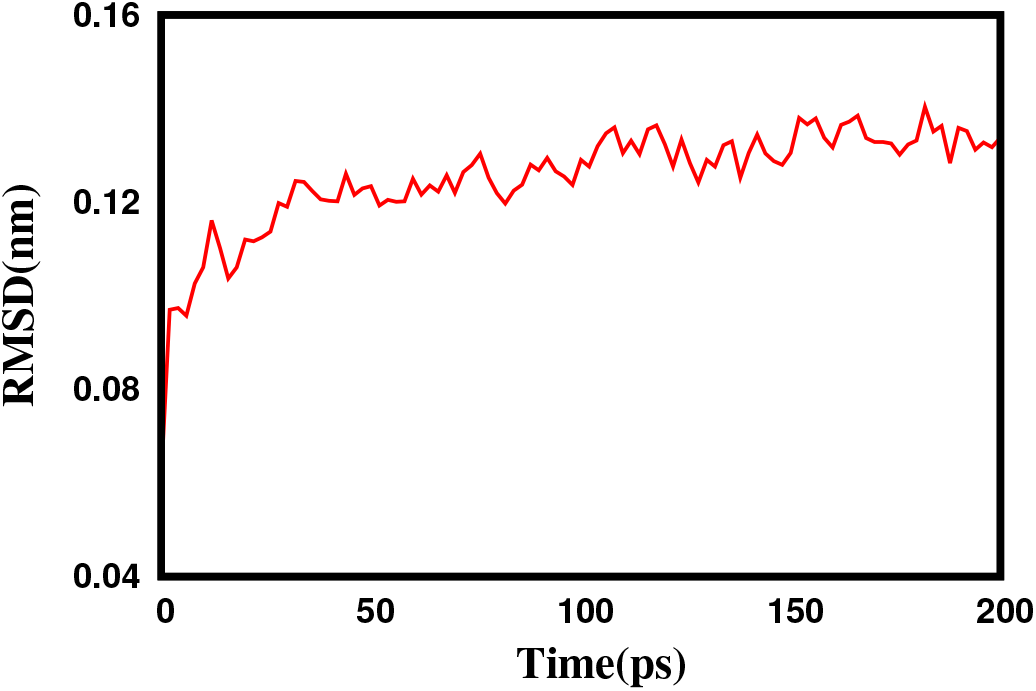
Root mean square displacement of all atoms fitted on the Protein_H (protein except hydrogens) group.

**FIG. 3.**
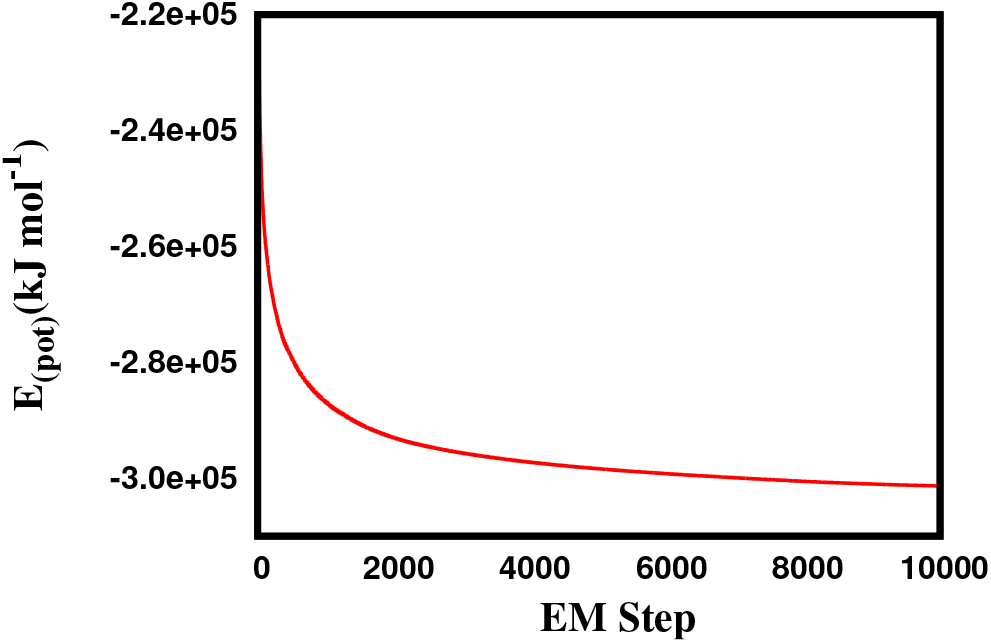
Average potential energy *Epot* as a function of energy minimization steps.

We also calculate the root mean square fluctuation (RMSF) which is a measure of the deviation between the position of *i*th-particle and some reference position with over the time T average. The root mean square fluctuation (RMSF) as a function of per residue over the `c_alpha` group is shown in fig. 4:(Upper Panel). The figure show the vibration around the equlibirum which depend on the local structure flexibility. The protein backbone is shown colour that vary with temperature. The red regions are at high fluctuation and the blue regions are relatively at low fluctuation.

**FIG. 4.**
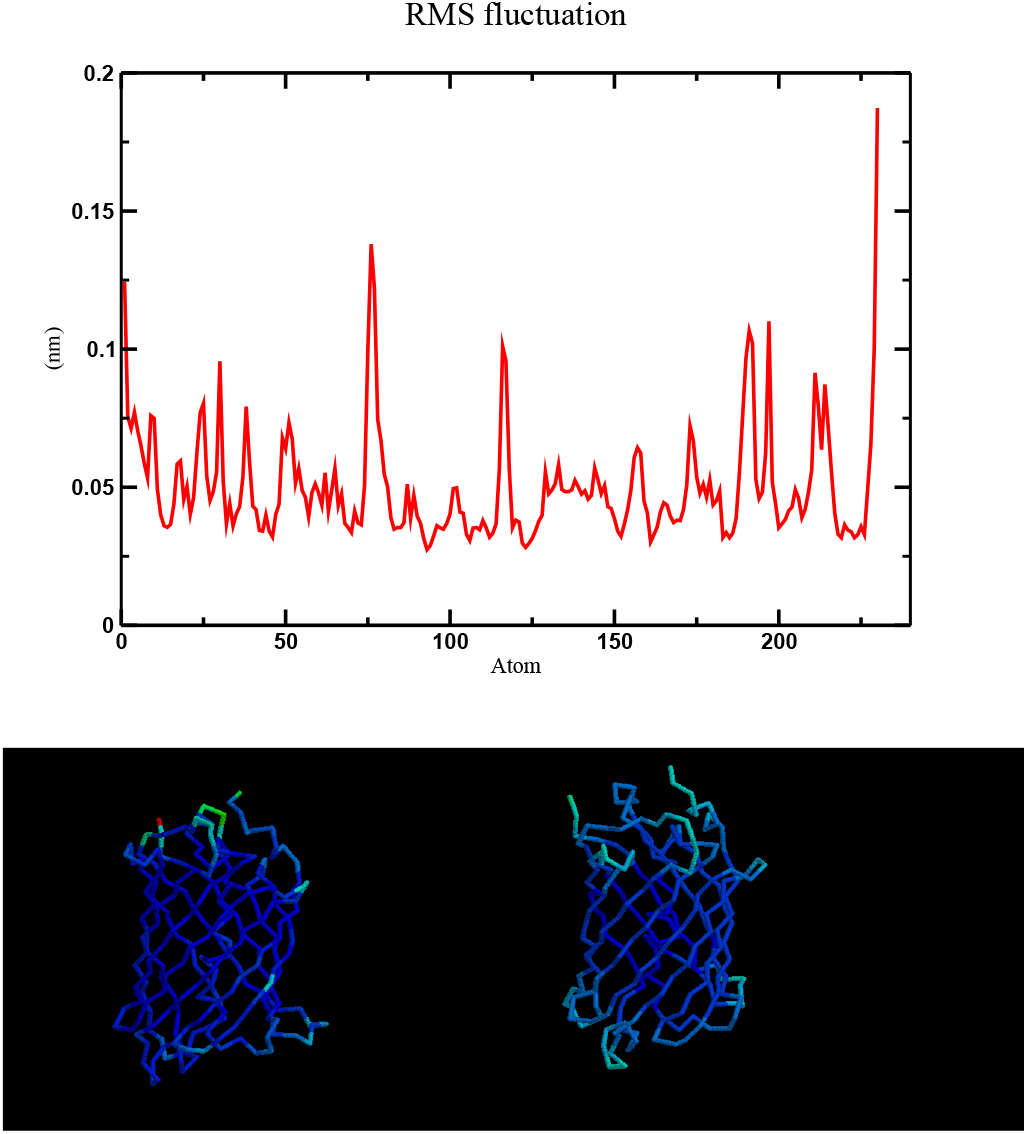
Upper Panel: Root mean square vale of fluctuation as a function of protein residue. Lower Panel: Backbone of Green fluorescent protein with color that vary according to temperature. The right panel with in lower panel show the comparison with the input GFP (PDB:1gfl.pdb). High fluctuation are represented in red color while the blue color represent the low fluctuation.

To further check the stability of the system, we plot the secondary structure of GFP per residue versus time using dssp (Dictionary of Secondary Structure for Proteins)-software, as shown in Fig. (5). The *β*-sheet and *α*-helix are highlighted with red and blue color respectively. Fig. (6) shows the Ramachandran plot of GFP. The figure describe the projection of the structure on the two di-hhedral angles *ϕ* (−180: 180) and Ψ (−180: 180) of the protein backbone. In Gromacs, `g_rama` command is implemented to obtained the Ramachandran plot.

**FIG. 5.**
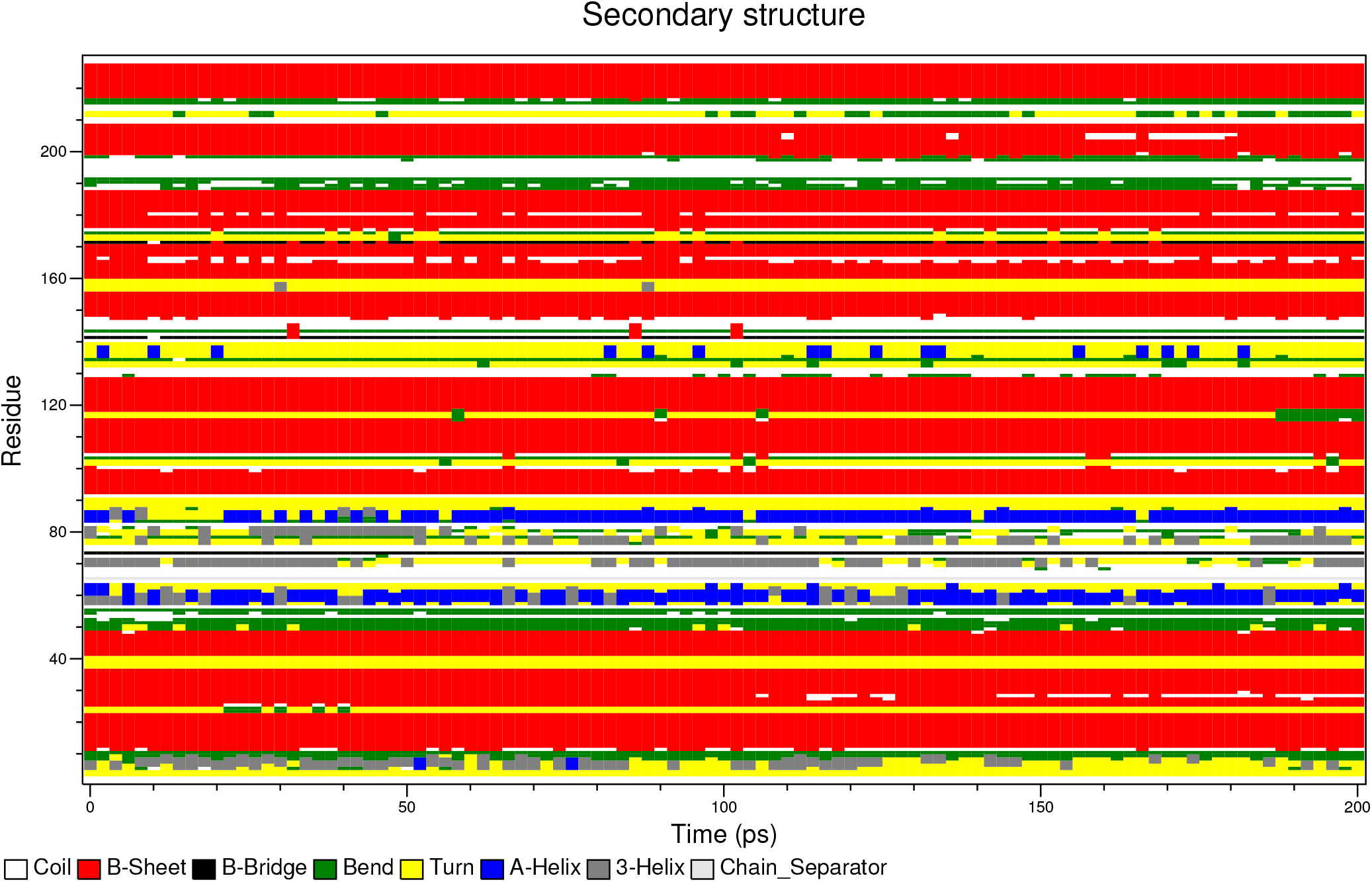
Secondary structure of GFP per residue versus time using dssp-software

**FIG. 6.**
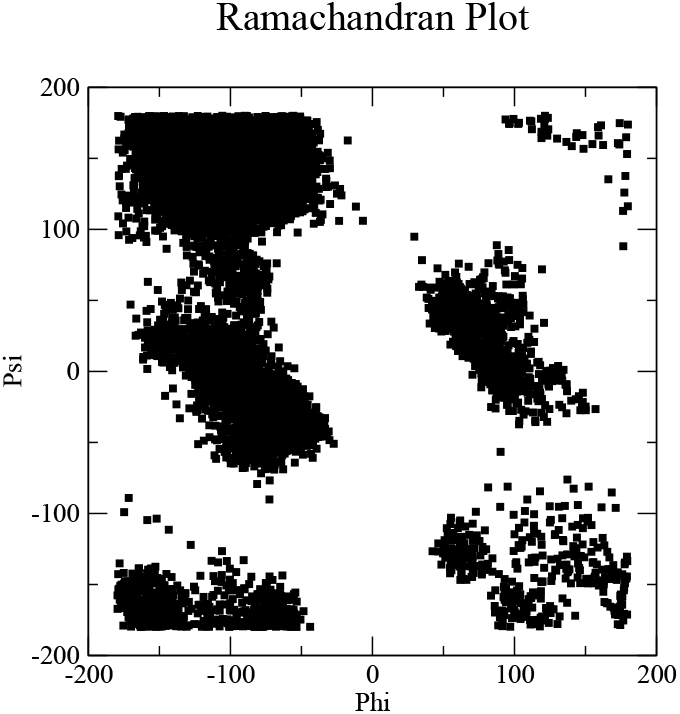
Ramachandran plot of a GFP.

## III. CONCLUSION

In summary, we discuss molecular dynamics simulation of GFP(Green Fluorescent Protein) with solvent water. We used steepest descent method to calculate the local energy minimum with 10^4^-steps. To determine whether the protein is close to the experimental structure during and after simulations, we calculate the root-mean-square deviation (RMSD) of the `Protein-H` atoms with respect to the X-ray structure. Secondary structure and Ramachandran plot of GFP protein is also presented.

